# High-throughput prediction of MHC Class I and Class II neoantigens with MHCnuggets

**DOI:** 10.1101/752469

**Authors:** XM Shao, R Bhattacharya, J Huang, IKA Sivakumar, C Tokheim, L Zheng, D Hirsch, B Kaminow, A Omdahl, M Bonsack, AB Riemer, VE Velculescu, V Anagnostou, KA Pagel, R Karchin

**Author notes:** These authors contributed equally (XM Shao, R Bhattacharya, J Huang, IKA Sivakumar). Corresponding author: Rachel Karchin, Ph.D, 217A Hackerman Hall, 3400 N. Charles St. Baltimore, MD USA 21204.

## Abstract

Computational prediction of binding between neoantigen peptides and major histocompatibility complex (MHC) proteins is an emerging biomarker for predicting patient response to cancer immunotherapy. Current neoantigen predictors focus on *in silico* estimation of MHC binding affinity and are limited by low positive predictive value for actual peptide presentation, inadequate support for rare MHC alleles and poor scalability to high-throughput data sets. To address these limitations, we developed MHCnuggets, a deep neural network method to predict peptide-MHC binding. MHCnuggets is the only method to handle binding prediction for common or rare alleles of MHC Class I or II, with a single neural network architecture. Using a long short-term memory network (LSTM), MHCnuggets accepts peptides of variable length and is capable of faster performance than other methods. When compared to methods that integrate binding affinity and HLAp data from mass spectrometry, MHCnuggets yields a fourfold increase in positive predictive value on independent MHC-bound peptide (HLAp) data. We applied MHCnuggets to 26 cancer types in TCGA, processing 26.3 million allele-peptide comparisons in under 2.3 hours, yielding 101,326 unique candidate immunogenic missense mutations (IMMs). Predicted-IMM hotspots occurred in 38 genes, including 24 driver genes. Predicted-IMM load was significantly associated with increased immune cell infiltration (p<2e−16) including CD8+ T cells. Notably, only 0.16% of predicted immunogenic missense mutations were observed in >2 patients, with 61.7% of these derived from driver mutations. Our results provide a new method for neoantigen prediction with high performance characteristics and demonstrate its utility in large data sets across human cancers.

**Synopsis:** We developed a new *in silico* predictor of Major Histocompatibility Complex (MHC) ligand binding and demonstrated its utility to assess potential neoantigens and immunogenic missense mutations (IMMs) in 6613 TCGA patients.

## Introduction

The presentation of peptides bound to major histocompatibility complex (MHC) proteins on the surface of antigen-presenting cells and subsequent recognition by T-cell receptors is fundamental to the mammalian adaptive immune system. Neoantigens derived from somatic mutations have been shown to be targets of immunoediting and drive therapeutic responses in cancer patients treated with immunotherapy (1,2). Because experimental characterization of neoantigens is both costly and time-consuming, many computational methods have been developed to predict peptide-MHC binding and subsequent immune response (3,4). Supervised neural network machine learning approaches are the best-performing (5–7) and the most widely used *in silico* methods. Despite these advances in computational approaches, improvements in predictive performance have been minimal, due in part to a lack of sufficiently large sets of experimentally characterized peptide binding affinities for most MHC alleles.

While neoantigen prediction for common MHC Class I alleles is well-studied (8), predictive accuracy on rare and less characterized MHC alleles remains poor (9,10) and Class II predictors are scarce(11). Current estimates suggest that Class II antigen lengths primarily range from 13-25 amino acids (12), and this diversity has been a major obstacle to developing *in silico* neoantigen predictors (11,13). As most neural network architectures are designed for fixed-length inputs, methods such as NetMHC (14–17) and MHCflurry (18) require pre-processing of peptide sequences or extensive training of separate classifiers for each peptide length.

Clinical application of MHC-peptide binding predictors, to identify biomarkers for cancer immunotherapy, requires scalability to large patient cohorts and low false positive rates (19). A cancer may contain hundreds of candidate somatically altered peptides, but few will actually bind to MHC proteins and elicit an immune response (20). For many years, most neoantigen predictors were trained primarily on quantitative peptide-HLA binding affinity data from *in vitro* experiments (21). Recent advances in immunopeptidomics technologies have enabled identification of thousands of naturally presented MHC bound peptides (HLAp) from cancer patient samples and cell lines (22) (19). Several new neoantigen predictors are trained only on HLAp data for Class I, for a limited number of peptide lengths (21) (23). The EDGE neural network is trained primarily on multi-allelic HLAp and RNAseq data from 74 cancer patients; ForestMHC is a random forest trained on HLAp from publicly available mono-allelic and deconvoluted multi-allelic cell lines. Furthermore, the potential to improve neoantigen predictors by integrating binding affinity and HLAp data (19) has motivated new hybrid approaches (14,18). However, most methods predict large numbers of peptides as candidate neoantigens, of which only a few are actually immunogenic in patients (11,19).

Here we present a long short-term memory (LSTM) neural network method, MHCnuggets, the first neoantigen predictor designed for MHC Class I and II alleles in a single framework. The method leverages transfer learning and allele clustering to accommodate both common, well-characterized MHC alleles and rare, less-studied alleles. While existing computational neoantigen predictors generate a large ranked list of candidate peptides, maximizing the number of highly-ranked true positives would be preferred in many applications (18). We demonstrate competitive predictive performance of MHCnuggets to widely-used methods on binding affinity datasets. In comparison to hybrid methods that have integrated binding affinity and HLAp data, we show decreased false positives and increased positive predictive value in a held-out cell line data set of ligands identified by mass spectrometry (7,24). To demonstrate the clinical utility and scalability of MHCnuggets to large patient cohorts, we investigated candidate immunogenic mutations from 26 tumor types in The Cancer Genome Atlas (TCGA). MHCnuggets yielded 101,326 candidate immunogenic missense mutations (out of 1,124,266) in less than 2.3 hours. These mutations were correlated with increased lymphocyte infiltration, however only 0.16% were observed in more than 2 patients.

## Methods

### Implementation

MHCnuggets uses a long short-term memory (LSTM) neural network architecture (25) (Figure 1A). LSTM architectures excel at handling variable length sequence inputs and can learn long-term dependencies between non-contiguous elements, enabling an input encoding that does not require peptide shortening or splitting (Figure 1B). LSTMs are capable of handling peptides of any length. In practice, a maximum peptide length should be selected for network training. We set maximum peptide input length of 15 for Class I and 30 for Class II, for computational efficiency purposes. These values cover the vast majority lengths observed in naturally presented MHC bound peptides (12). The networks were trained with transfer learning (26), which allows networks for less well-characterized alleles to leverage information from extensively studied alleles (Figure 1C). Transfer learning was also used to train networks combining binding affinity and HLAp datasets. In addition, MHCnuggets’ architectures can be trained using either continuous binding affinity measurements from *in vitro* experiments (half maximal affinity or IC50) and/or immunopeptidomic (HLAp) binary labels. The former utilizes a mean-squared error (MSE) loss while the latter utilizes binary cross-entropy (BCE) loss for training.

**Figure 1.**
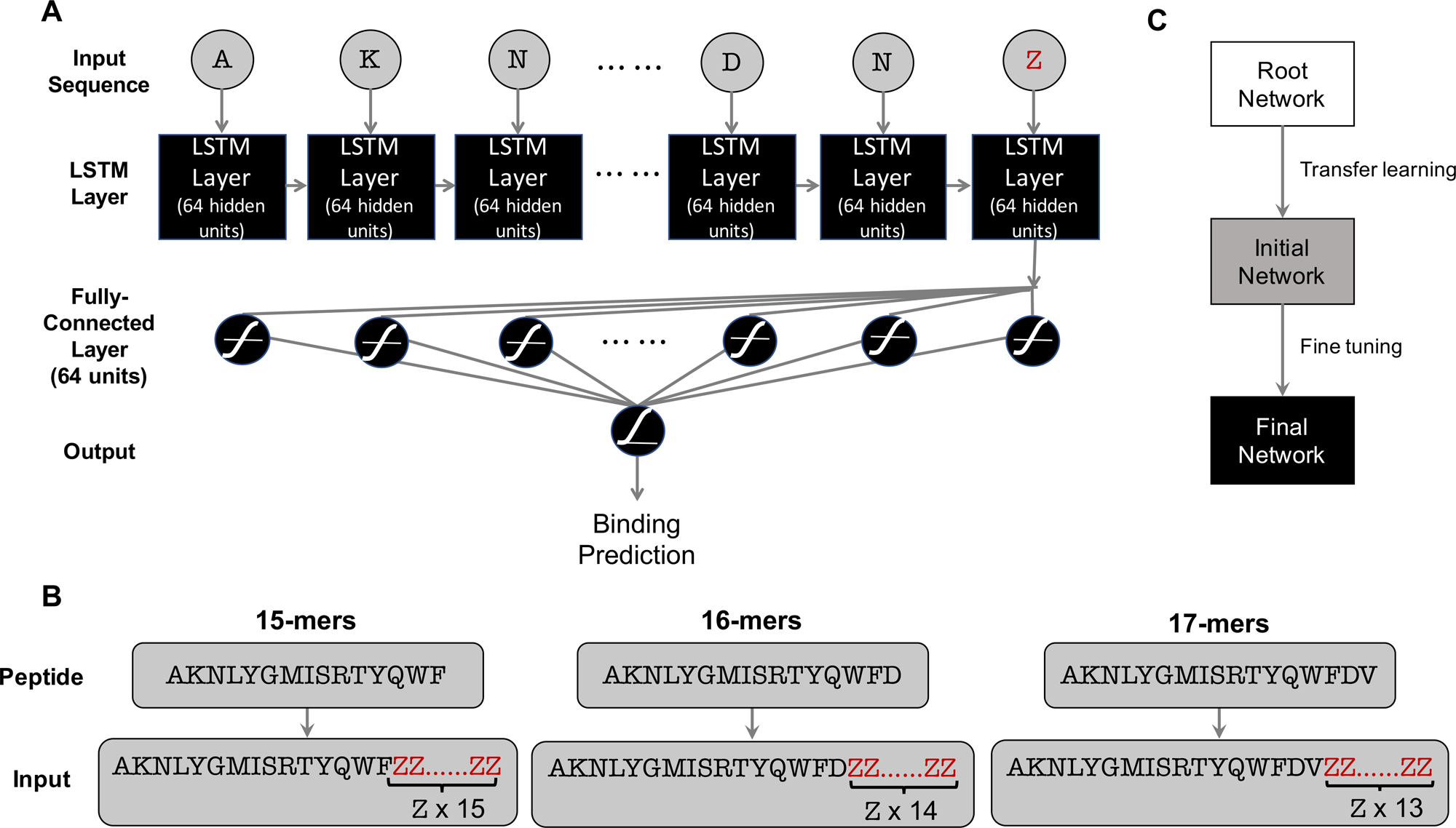
**A) MHCnuggets’ architecture.** A network is trained for each MHC allele. Each network has a LSTM layer with 64 hidden units, a Fully Connected (FC) layer with 64 hidden units and a final output layer of a single sigmoid unit. **B) Input scheme for peptides with variable lengths.** MHCnuggets architecture is capable of handling peptides of any length, but in practice a maximum length should be selected. Peptides are extended with padding until they reach the maximum length, prior to input into the neural network. The example shows padding for Class II peptides with maximum length set to 30 amino acids. **C) Transfer learning protocol for parameter sharing among alleles.** A base allele-specific network is trained for each MHC class, with an allele selected by largest number of training examples. Transfer learning is applied to train networks for the remaining alleles with initial network weights set to final base network weights. A fine tuning step identifies alleles that can be leveraged for a second round of transfer learning to produce a final network (detail in Supplementary Methods).

For each MHC allele, we trained a neural network model consisting of an LSTM layer of 64 hidden units, a fully connected layer of 64 hidden units and a final output layer of a single sigmoid unit (Figure 1A). Specifics of peptide encoding, training loss functions, optimization and regularization methods are in Supplementary Methods. For the 16 alleles where allele-specific HLAp training data was available (27), we trained networks on both binding affinity and HLAp data (MHCnuggets). Next, we trained networks only with binding affinity measurements (MHCnuggets noMS) for all MHC Class I alleles. Due to the lack of allelic-specific HLAp training data for Class II, all MHC Class II networks were trained only on binding affinity measurements. In total, we trained 148 Class I and 136 Class II allele-specific networks. Common alleles comprise a small fraction of all known MHC alleles (28). To handle binding predictions for rare alleles, MHCnuggets selects a network by searching for the closest allele, based on previously published supertype clustering approaches. We prioritized approaches based on binding pocket biochemical similarity when available. Briefly, HLA-A and HLA-B alleles were clustered by MHC binding pocket amino acid residue composition (29), and HLA-C and all MHC II alleles were hierarchically clustered based upon experimental mass spectrometry and binding assay results (30,31). For alleles with no supertype classification, the closest allele was from the same HLA gene, and allele group if available, with preference for alleles with the largest number of characterized binding peptides. All networks were implemented with the Keras Python package (TensorFlow back-end) (32,33). Open source software is available at https://github.com/KarchinLab/mhcnuggets, installable via pip or Docker, and has been integrated into the PepVacSeq (34), pvactools (35) and Neoepiscope (36) pipelines.

### Benchmarks

To accurately assess the performance of MHCnuggets on a variety of MHC-peptide binding prediction tasks, we utilized six distinct benchmark sets: MHC Class I alleles, MHC Class II alleles, common alleles with a trained model (allele-specific prediction) and rare alleles (pan-allele prediction) (Figure 2, Table S1). To compare to the widely-used HLA ligand prediction tools from the NetMHC group (NetMHC3.0, NetMHC 4.0, NetMHCpan2.0, NetMHCpan 4.0) (16,17), which can be trained only by their developers, as well as the open source MHCflurry tools, we employed multiple benchmarking strategies: 1) independent benchmark test set of peptides not included as training data for any of the methods; 2) a previously published paired training/testing benchmark; 3) five-fold cross-validation benchmark; 4) leave-one-molecule-out (LOMO) benchmark.

**Figure 2.**
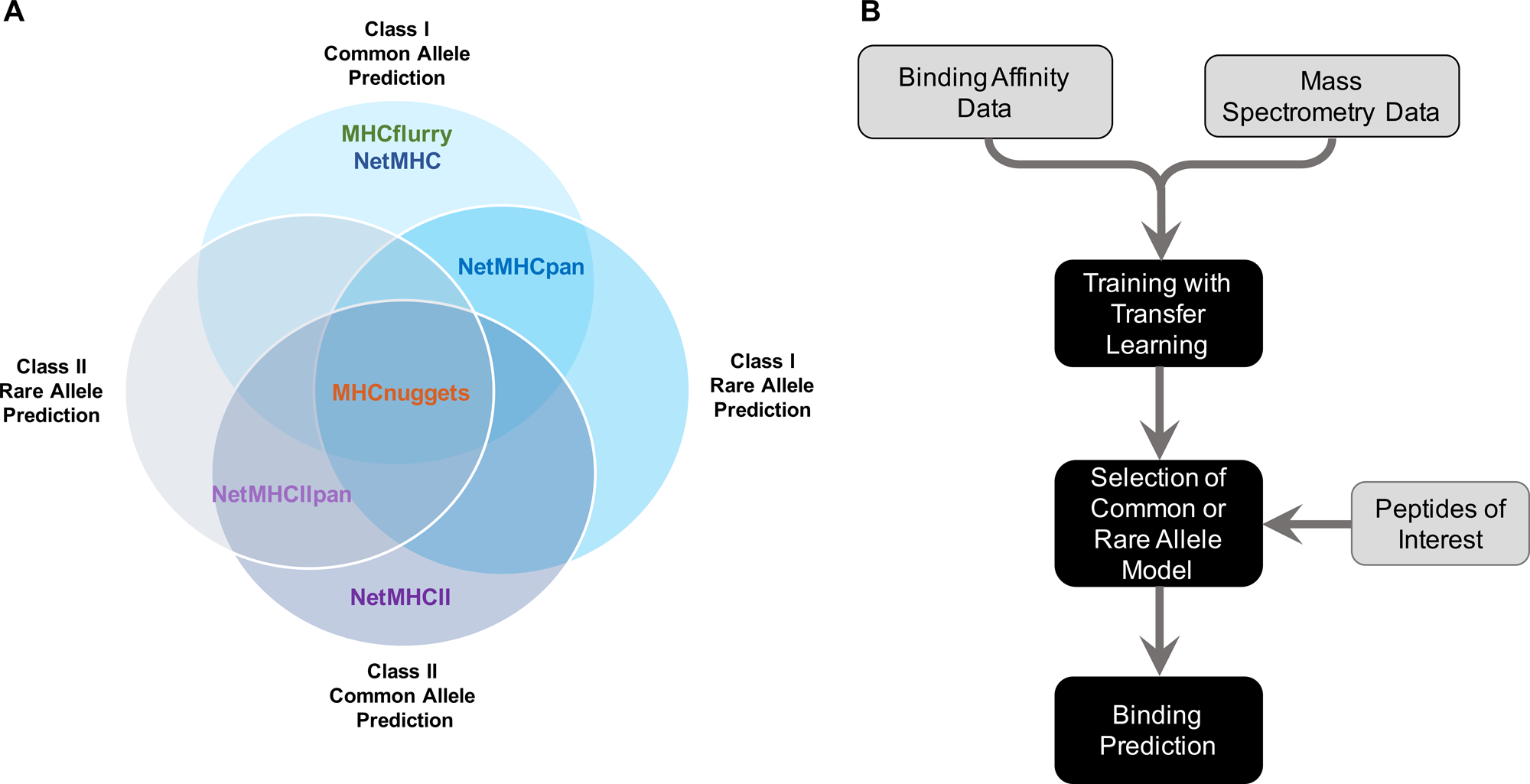
MHCnuggets’ features. A) Venn diagram representation of the MHC-peptide binding prediction functions of MHCnuggets and several other currently available tools. B) Training and MHC allele model selection scheme for MHCnugget**s.**

We evaluated six MHC Class I predictors on independent binding affinity and HLAp datasets (7,8,24). First, we compared MHCnuggets to several Class I predictors that incorporate both binding affinity and HLAp data: MHCflurry 1.2.0, MHCflurry (train-MS), NetMHC 4.0, and NetMHCpan 4.0. Each method was benchmarked using an independent set of MHC-bound peptides identified by mass spectrometry across seven cell-lines for six MHC I alleles (Bassani-Sternberg 2017, Trolle 2016). For testing, HLAp hits were combined with random decoy peptides sampled from the human proteome in a 1:999 hit-decoy ratio, as described by Abelin *et al.* (27), totaling 23,971,000 peptides. Next, four MHC Class I predictors trained only on binding affinity data (MHCnuggets (noMS) and MHCflurry (noMS), NetMHC 3.0 and NetMHCpan 2.0) were evaluated with the Kim *et al.* dataset (5), in which each predictor was trained with the BD2009 data and tested on BLIND data. It was possible to compare NetMHC3.0 and NetMHCpan2.0 performance on Kim et al., because they have previously published predicted IC50s for all peptide-MHC pairs in BLIND. This allowed us to calculate their PPV_n_, area under the ROC curve (auROC), Kendall’s *tau*, and Pearson’s *r* correlations.

Next, we compared MHCnuggets’ Class II ligand prediction performance with self-reported performance statistics of NetMHC group’s MHC Class II methods (37). We used the Jensen *et al*. five-fold cross-validation benchmark to assess allele-specific MHC Class II prediction of MHCnuggets and NetMHCII 2.3, for 27 alleles. NetMHCII 2.3 reported the average auROC for five-fold cross-validation, and we report MHCnugget’s positive predictive value for each of the 27 alleles as well as the average auROC, Pearson’s *r* and Kendall-Tau correlations.

The leave-one-molecule-out (LOMO) benchmarks are a type of cross-validation designed to estimate the performance of peptide binding prediction with respect to rare MHC alleles. Given training data for *n* MHC alleles, the data for a single allele is held out and networks are trained for the remaining *n−1* alleles. Then for each peptide, predictions are generated by the remaining networks. We designed a LOMO benchmark to evaluate MHC Class I rare allele prediction, by selecting 20 alleles with 30 to 100 characterized peptides in IEDB. For Class II rare allele prediction, we used the Jensen et al. LOMO benchmark. We were unable to assess rare allele prediction for NetMHC Class I methods, as no published results were available. For the NetMHC Class II methods, we compared MHCnuggets to their self-reported auROCs.

### TCGA analysis pipeline

To assess candidate immunogenic somatic mutations in patients from the TCGA cohort, we developed and implemented a basic pipeline based on whole-exome and RNA sequencing data. Our analysis builds upon work from the TCGA PanCancer Analysis teams for drivers (38), mutation calling (39) and cancer immune landscapes (40). We obtained somatic mutation calls for all cancer types from Multi-Center Mutation Calling in Multiple Cancers (MC3) (v0.2.8) (7775 patients). Tumor-specific RNA expression values from Broad TCGA Firehose were standardized across tumor types using the RSEM Z-score (41). MHC allele calls were obtained from the TCGA cancer immune landscape publication, in which up to six MHC Class I alleles (HLA-A, HLA-B, and HLA-C) were identified for each patient using OptiType (42). We included patients for which mutation calls, MHC allele calls and RNA expression values were available from TCGA (Supplementary Methods). After these considerations, the analysis included 6613 patients from 26 TCGA tumor types. Six cancer types were not included in our analysis, because 15 or fewer patients met this requirement: Lymphoid Neoplasm Diffuse Large B-cell Lymphoma (DLBC), Esophageal carcinoma (ESCA), Mesothelioma (MESO), Skin Cutaneous Melanoma (SKCM), Stomach adenocarcinoma (STAD), Ovarian serous cystadenocarcinoma (OV).

The somatic missense mutations identified in each patient were filtered to include only those with strong evidence of mutant gene RNA expression in that patient (Z>=1.0). For each mutation that passed this filter, we used the transcript assigned by MC3 to pull flanking amino acid residues from the SwissProt database (43), yielding a 21 amino acid residue sequence fragment centered at the mutated residue. All candidate peptides of length 8,9,10 and 11 that included the mutated residue were extracted from each sequence fragment. Next binding affinity predictions were generated for each mutated peptide for up to six MHC Class I alleles, depending on the patient’s HLA genotypes. In total, each somatic mutation was represented by 38 mutated peptides for up to 6 possible MHC pairings.

We applied a permissive filter to select candidate immunogenic peptides, requiring mutated peptides to have binding affinity of IC50<500nM for at least one MHC allele. Somatic missense mutations that generated neoantigens meeting these criteria were considered candidate immunogenic missense mutations (IMMs). For a given patient, if a mutation was predicted to be a candidate IMM for multiple alleles, it was counted only once using the MHC allele with the lowest predicted IC50. Finally, for each patient we counted the number of predicted IMMs found in their exome and stratified by tumor type. We then identified predicted IMMs that were harbored by more than one patient.

We sought to ascertain whether predicted IMMs occurred preferentially in particular gene or protein regions. Using the HotMaps 1D algorithm v1.2.2 (46), we clustered primary amino acid residue sequence to identify regions where mutations were frequently predicted as IMM, with statistical significance (q<0.01, Benjamini-Hochberg method (47)). In this analysis, mutations were stratified by cancer type, and we considered enrichment within linear regions of 50 amino acid residues.

We considered that mutation immunogenicity might be associated with potential driver status of a mutation. Driver status was inferred by CHASMplus (48), a random forest classifier that utilizes a multi-faceted feature set to predict driver missense mutations. It has been previously shown to be effective at identifying both common and rare driver mutations. For each mutation, its immunogenicity was represented as a binary response variable and driver status was used as a covariate. Mutations with CHASMplus q-value < 0.01 were considered drivers (48). We modeled the relationship with univariate logistic regression (R glm package with binomial link logit function).

To assess whether the total number of predicted IMMs per patient was associated with changes in tumor immune infiltrates, we performed Poisson regression (R glm package with Poisson link log function). All estimates of immune infiltrates were obtained from Thorsson et al. (40,49). We fit two univariate models in which the response variable was the predicted IMM count and the covariate was either total leukocyte fraction or fraction of CD8+ T-cells.

## Results

### High-throughput MHCnuggets breaks the MHC ligand prediction plateau

The MHCnuggets LSTM neural network architecture accepts peptides of variable lengths as inputs so that ligand binding prediction can be performed for both MHC Class I and Class II alleles. To enable binding prediction for rare MHC alleles with limited experimental data, we designed a method that leverages networks built for closely-related common alleles with extensive data. When available, we utilize a transfer learning protocol to integrate binding affinity and HLAp results in a single network model, to better represent the natural diversity of MHC-binding peptides.

To assess the baseline performance assessment for MHCnuggets’ allele-specific networks on binding affinity data, we compared our approach with the most widely used MHC Class I ligand prediction methods, using two validation sets of binding affinity measurements (Kim et al. (5) Bonsack et al. (8)). We trained and tested MHCnuggets (noMS) and MHCflurry (noMS) using the Kim *et al* dataset, and evaluated the predictions provided by NetMHC 3.0 and NetMHCpan 2.0. We observed that MHCnuggets’ performance (PPV_n_ = 0.829, auROC=0.924) was comparable to these methods (Figure 3a) (PPV_n_ of all methods=0.825 +/− 0.005, auROC of all methods = 0.928 +/− 0.0031). MHCnuggets was also comparable (PPV_n_ = 0.633, auROC=0.794) to these methods when tested on the Bonsack *et al*. dataset (PPV_n_ of all methods = 0.625 +/− 0.008, auROC of all methods = 0.77 +/− 0.02) (Figure 3A) (+/− refers to standard deviation) (Table S3a, S3b, Table S4a, S4b).

**Figure 3.**
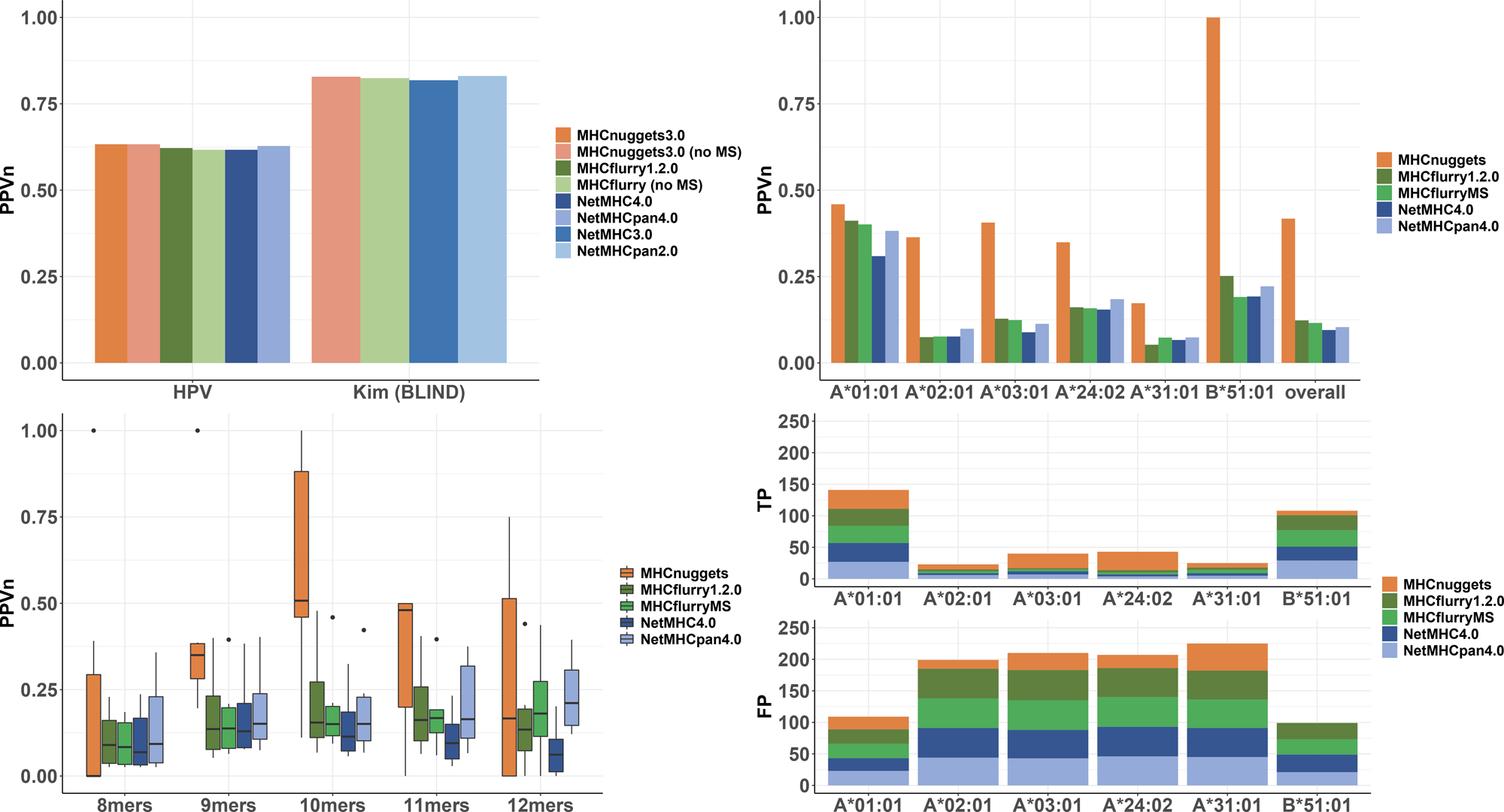
MHC Class I benchmark comparisons. A) PPV_n_ for MHC Class I allele-specific prediction on binding affinity test sets from Bonsack et al. (7 alleles) and Kim *et al*. (53 alleles) B) PPV_n_ for MHC Class I allele-specific prediction on HLAp BST data set (Bassani-Sternberg et al. and Trolle *et al*.), stratified by allele (6 alleles). C) PPV_n_ for MHC Class I allele-specific prediction on HLAp BST data set (from B) stratified by peptide sequence length. D) True and false positives for each method on the top 50 ranked peptides from the HLAp BST data set. PPV_n_ = positive predictive value on the top *n* ranked peptides, where *n* is the number of true binders. TP=true positives. FP=false positives.

Historically, neoantigen prediction methods have focused on Class I and trained on binding affinity data from IEDB (50). More recent work has incorporated both binding affinity and HLAp data into network training (14,18). We compared MHCnuggets with several Class I predictors that also used both binding affinity and HLAp data: MHCflurry 1.2.0, MHCflurry (train-MS), NetMHC 4.0, and NetMHCpan 4.0. We selected the Bassani-Sternberg/Trolle (BST) HLAp dataset (7,24,27) as an independent benchmark, as it was not previously included as training data by any of these methods. For all alleles tested, MHCnuggets achieved an overall PPV_n_ of 0.42 and auROC of 0.82 (Figure 3B). On average, MHCnuggets’ PPV_n_ was more than three times higher than MHCflurry 1.2.0, MHCflurry (train-MS), NetMHC 4.0, and NetMHCpan 4.0. For all alleles, MHCnuggets predicted substantially fewer binders than other methods, resulting in fewer false positive predictions. Stratifying by peptide length, MHCnuggets’ increased PPV_n_ was most prominent for peptides of length 9, 10, and 11 (Figure 3C). The length distribution of predicted binders was also commensurate with the observed distribution of naturally occurring binders in the HLAp benchmark tests (Trolle 2016; Table S5a, S5b, S5c, S5d).

For some clinical applications, it may be desirable to minimize the number of false positives among a small number of top-scored peptides. We also compared PPV of the methods listed above on their top 50 and 500 ranked peptides from the BST dataset (six MHC Class I alleles). MHCnuggets exhibited the highest PPV in the top 50 for all alleles except HLA-B*51:01 and the highest PPV in the top 500 for all alleles (Figure 3D, Table S5e).

### Prediction of peptide-MHC binding for Class II and rare alleles

We assessed baseline performance of MHCnuggets Class II allele-specific networks on binding affinity data. To enable comparison with the Class II methods from the NetMHC group, we used a five-fold cross validation benchmark derived from IEDB that was included in the publication describing NetMHCII-2.3 and NetMHCIIpan-3.2 (37). First, we computed PPV_n_ for each of the 27 allele-specific networks separately (Figure 4A) (mean PPV_n_=0.739). Next, we computed the overall auROC, Pearson r and Kendall Tau correlations for all 27 Class II alleles. MHCnuggets overall auROC (0.849) was comparable to that of the NetMHCII-2.3 (0.861) and NetMHCIIpan-3.2 (0.861). Comparison to NetMHC Class II methods was limited to overall auROC as published in (37), because their results are not publicly available (Figure 4B) (Table S6a, Table S6b).

**Figure 4.**
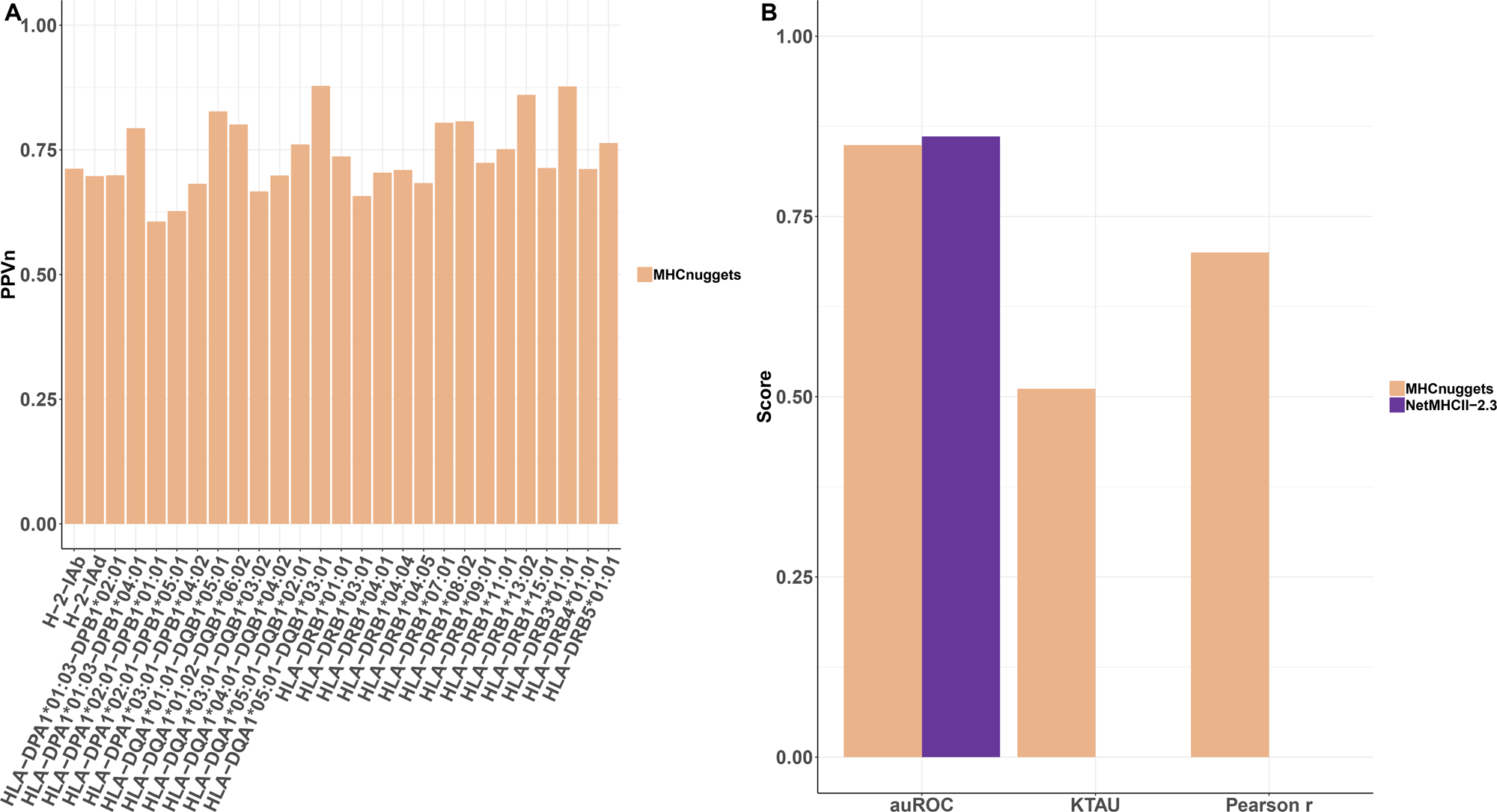
MHC Class II benchmark comparisons. A) PPV_n_ for MHC Class II allele-specific prediction on binding affinity test set from Jensen *et al.* (27 alleles, stratified by allele). B) auROC, K-Tau, Pearson r scores for MHC Class II alleles from five-fold cross-validation. NetMHCII2.3 performance is from their self-reported auROC. auROC= area under the receiving operator characteristic curve. K-Tau = Kendall’s *tau* correlation. PPV_n_ = positive predictive value on the top *n* ranked peptides, where *n* is the number of true binders.

We estimated performance for those Class I and Class II MHC alleles for which we were unable to train allele-specific networks, using leave-one-molecule-out (LOMO) cross-validation (37). In this LOMO protocol, MHC-peptide binding is assessed for a well-characterized allele that has been held out from training, to approximate prediction performance for a rare allele (Figure 5A). For the 20 Class I alleles, the mean PPV_n_ was 0.65 and the mean auROC was 0.671. For the 27 Class II alleles, the mean PPV_n_ was 0.65 and the mean auROC was 0.792. In comparison, the Class II mean auROC of NetMHCIIpan-3.2 was 0.781 (Figure 5B, Figure 5C). Further performance results of NetMHCpan rare allele predictors for both Class I and Class II were not publicly available for LOMO tests (Table S7, Table S8a, Table S8b).

**Figure 5.**
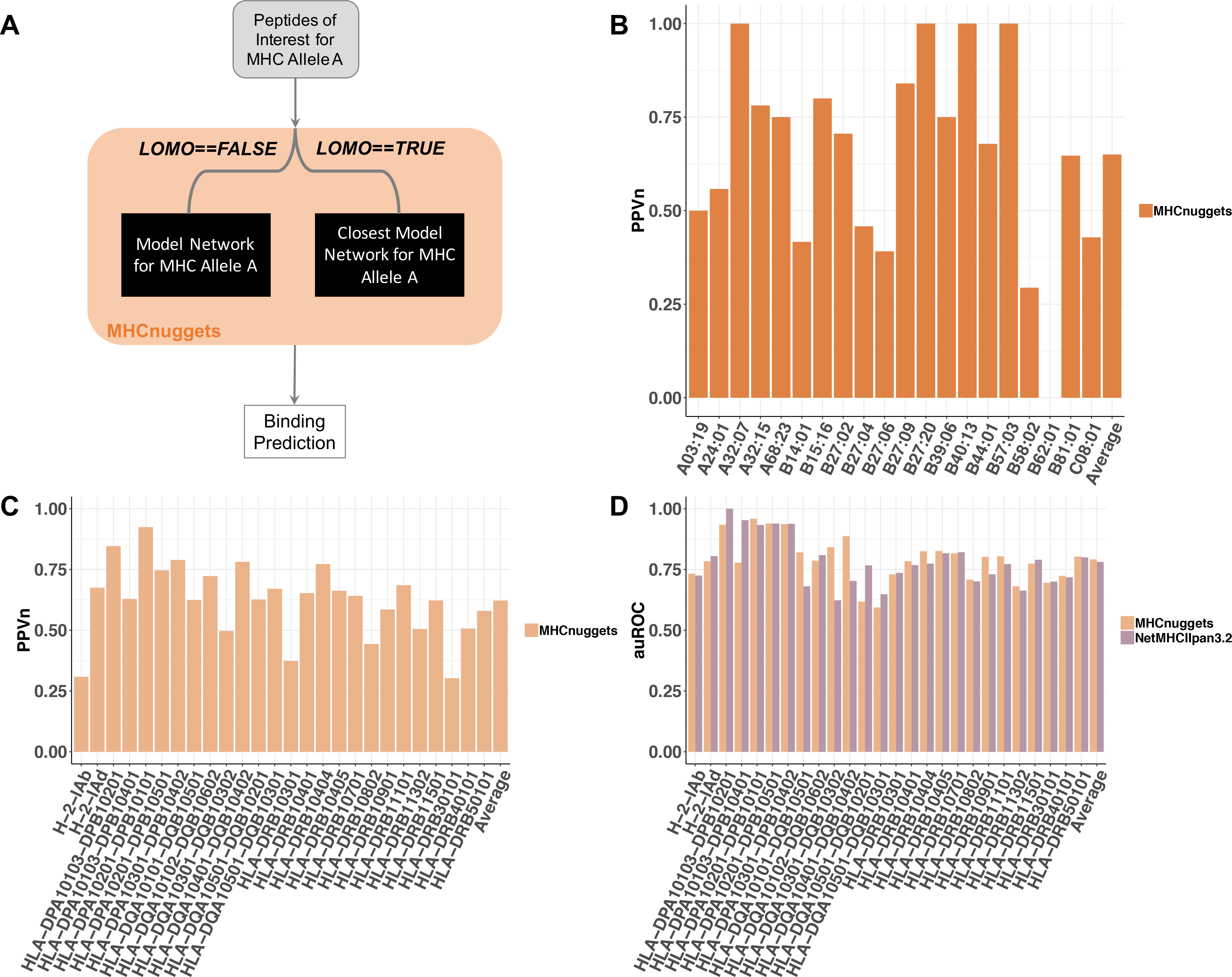
MHC Class I and II benchmark comparisons to estimate rare allele performance. A) Schematic representation of leave one molecule out (LOMO) testing. B) PPV_n_ for MHC Class I rare allele prediction on IEDB pseudo-rare alleles binding affinity test set (20 alleles, stratified by allele). C) PPV_n_ for MHC Class II rare allele prediction on binding affinity test set from Jensen et al. (27 alleles, stratified by allele). D) auROC for MHC Class II rare allele prediction on LOMO binding affinity test set from Jensen *et al*. (27 alleles, stratified by allele). NetMHCIIpan3.2 results are from their self-reported auROC. auROC = area under the receiving operator characteristic curve. PPV_n_ = positive predictive value on the top *n* ranked peptides, where *n* is the number of true binders.

### Fast and scalable computation

When run on a GPU architecture, MHCnuggets was substantially faster and scaled more efficiently than MHC ligand predictors from the NetMHC family and MHCflurry. Given an input of one million peptides randomly selected from Abelin et al., MHCnuggets runtime was 4.5, 3.2, and 18 times faster than MHCflurry 1.2.0, NetMHC 4.0, NetMHCpan 4.0, respectively (Figure 6A). The improvement was even more pronounced for Class II peptides, for which an input of one million peptides to MHCnuggets ran 65.6 times and 126 times faster than NetMHCII2.3 and NetMHCIIpan 3.2, respectively (Figure 6B). As the total number of input peptides was increased from 0 to one million, the runtime per peptide plateaued for other methods but decreased exponentially for MHCnuggets.

**Figure 6.**
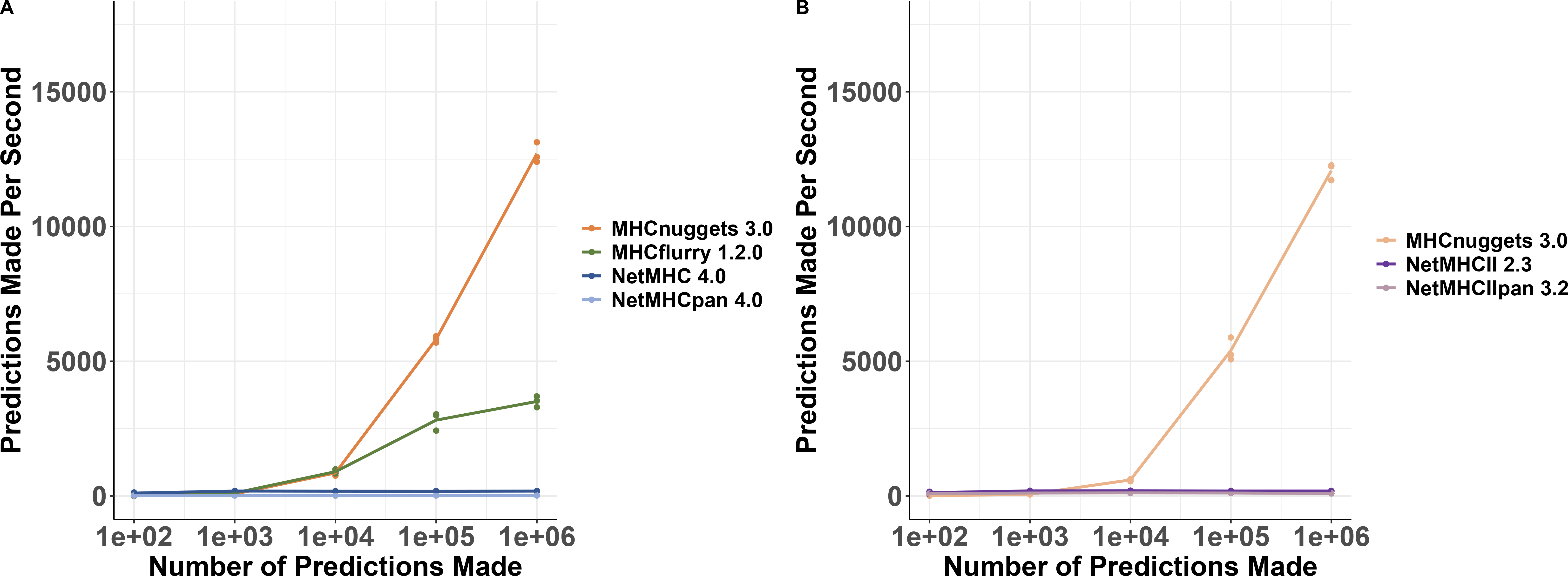
Timing and scalability. Runtime benchmark of most recent version of tested methods over a range of inputs (up to 1 million peptides). A) MHC Class I prediction. B) MHC Class II prediction

### Candidate MHC Class I immunogenic missense mutations in TCGA patients

To illustrate the utility of MHCnuggets’ improvements in scalability and positive predictive value for the analysis of very large patient cohorts, we predicted Class I *immunogenic missense mutations* (IMMs) in patients sequenced by the TCGA consortium (Methods). In our analysis pipeline, patient exomes were split into 21 amino acid residue sequence fragments, centered on each somatic missense mutation. For each sequence fragment, MHCnuggets predicted the MHC binding for all possible 8-, 9-, 10- and 11-length peptide windows. Peptides that passed filters of predicted IC50 threshold (<500nM) and gene expression (Z>1.0) (Methods) for at least one patient-specific MHC allele were classified as predicted IMMs (Table S9a). Finally, we characterized driver status and positional hotspot propensity of the predicted IMMs.

Total processing time for 26,284,638 allele-peptide comparisons supported by RNAseq expression was under 2.3 hours. First, we sought to ascertain the extent of variability in predicted IMM count among individuals with different cancer types. Next, we identified predicted IMMs and protein regions enriched for predicted IMMs that were shared across patients, because these might be informative for neoantigen-based therapeutic applications. Then we considered whether predicted IMMs were more or less likely to be driver mutations. Finally, we assessed the associations between predicted patient IMM load and computationally estimated immune cell infiltrates.

After applying a strict gene expression filter, we identified 101,326 unique predicted IMMs in 26 TCGA cancer types, with a mean of 15.6 per patient. We found that the majority of patients harbored fewer than 6 predicted IMMs, and 197 patients had none. Seventy-two percent of patients had from 1 and 10 predicted IMMs, compared to 1.9% of patients with more than 100, and nine patients with more than 1000 (Figure 7A). Cancer types with the highest number of predicted IMMs were uterine corpus endometrial carcinoma (UCEC), colon adenocarcinoma (COAD), and lung adenocarcinoma (LUAD), previously known for high mutation burden and immunogenicity (40). UCEC and COAD are also known to have a high frequency of microsatellite-instable (MSI) tumors. The lowest number were found in Uveal Melanoma (UVM), Paraganglioma & Pheochromocytoma (PCPG), and Testicular Germ Cell Cancer (TGCT) (Figure 7B, Table S9b).

**Figure 7.**
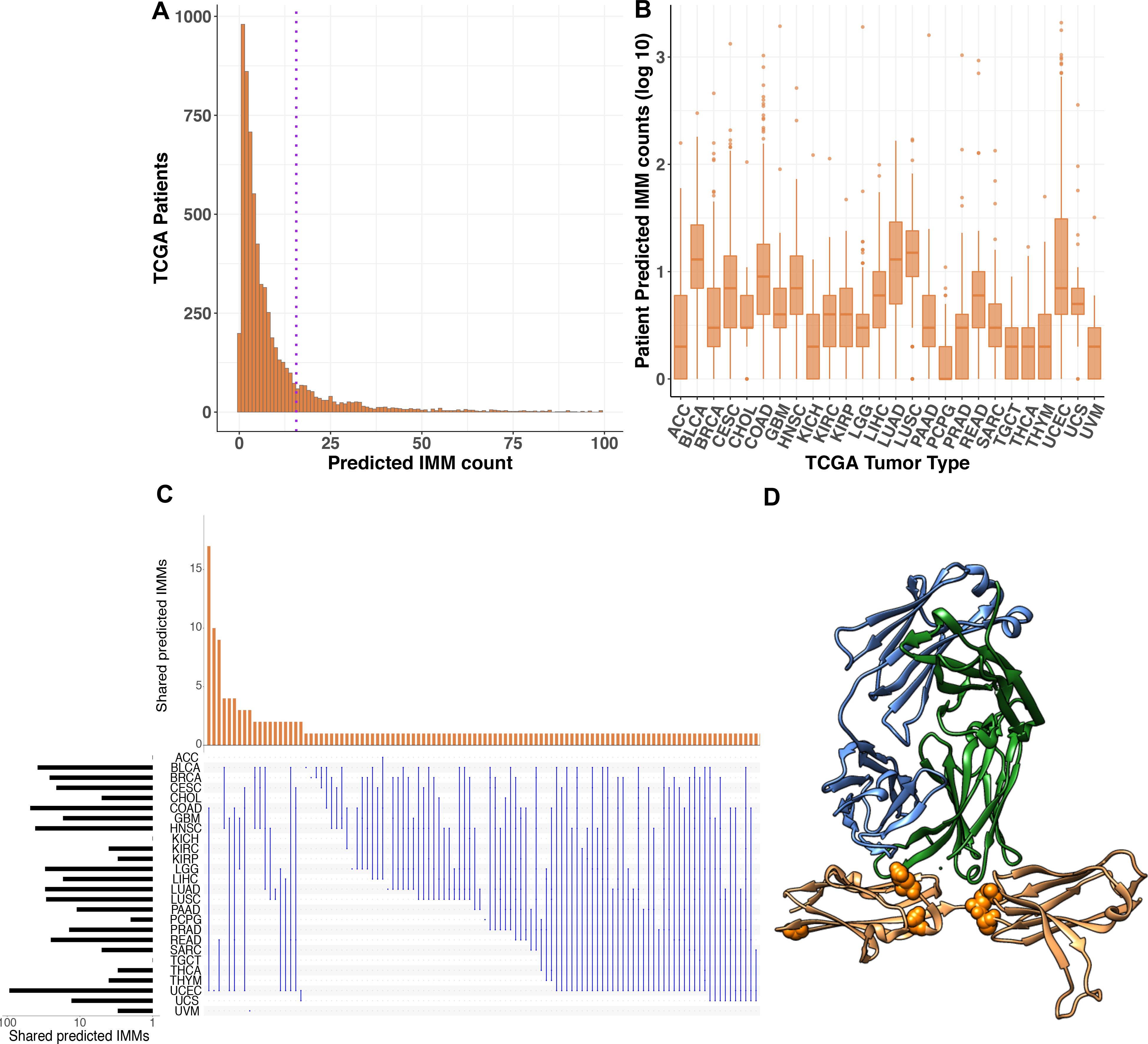
MHC Class I IMMs in TCGA patients. A). Number of predicted immunogenic missense mutations (IMMs) identified in 6,613 TCGA patients. Dotted line = mean IMMs per patient (15.6). Note, 123 patients had >100 predicted IMMs but are not included for visual clarity. B) Number of predicted IMMs by cancer type. C) IMMs shared by three or more patients and the cancer types in which they occurred. Each row represents a cancer type and each column illustrates the overlap of IMMs seen in a single cancer type or multiple cancer types. For example, the first column shows the number of IMMs shared among patients with colorectal adenocarcinoma (COAD) and uterine corpus endometrial carcinoma (UCEC). Bars to the left show the total number of unique IMMs in each cancer type. *Bar heights are count of unique shared IMMs, not total number of patients in which the IMM was observed. Cancer type abbreviations are in Supplementary Methods. Image generated with UpSetR (58). D) Fibroblast growth factor receptor (*FGFR3*) IMM hot region identified by HotMAPs in bladder cancer (BLCA). IMMs shown and number of BLCA patients with the IMM: p.E216K (1), p.D222N (1), p.G235D (1) p.R248C (3) and p.S249C (24). Except for p.G235D, these IMMs are proximal to the interface of FGFR3 protein and the light and heavy chains of an antibody fragment designed for therapeutic application in bladder cancer (PDB ID: 3GRW) (59).

Across all cancer types, we identified 1,393 predicted IMMs harbored by two or more patients, of which 167 were identified in three or more patients. Of these 167 only 11.5% occurred exclusively in a single cancer type (Figure 7C). The predicted IMMs identified in the largest number of patients were *IDH1* R132H (62), *FGFR3* S249C (24), *PIK3CA* E545K (23), KRAS G12D (18), *PIK3CA* E542K (18), TP53 R175H (18), TP53 R248Q (18), *TP53 R273C (17)*, and *KRAS* G12V (16), which are well known recurrent oncogenic driver mutations (51,52). Of the 1071 genes harboring predicted IMMs in 2 or more patients, the ones containing the most included TP53 (68), CTNNB1 (18), PIK3CA (16), HRAS (8), KRAS (7), PTEN (7), FBXW7 (6), EGFR (5), MDN1 (5), POLE (5), TRRAP (5), and VPS13C (5) (Table S9c). Six missense mutations harbored by patients in the TCGA cohorts were previously validated by CD8+ T cell response assays (53) (54) (55). Of the six missense mutations, *TP53*^*R248Q*^*, TP53*^*Y220C*^*, TP53*^*R175H*^*, TP53*^*R248W*^ *and KRAS*^*G12D*^ were predicted to be IMMs by our MHCnuggets pipeline and were shared by three or more of the TCGA patients.

Furthermore, 61.7% of the 167 predicted IMMs shared by three or more patients were classified as driver missense mutations by CHASMplus (q<0.01). This percentage is significantly higher than the number of predicted drivers among all TCGA missense mutations (9,821 out of 791,637 or 1.2%). It is worth noting that while many shared IMMs were predicted to be driver missense mutations, the percentage of predicted IMMs predicted to be drivers was ~0.1% of total predicted IMMs in our study. When compared to the OncoKB database of experimentally confirmed driver mutations (56), 53.9% of the shared predicted IMMs identified as “oncogenic” or “likely oncogenic” driver mutations. The percentage is lower (25.7%) if “likely oncogenic” mutations are excluded.

While we observed a limited number of shared IMMs, we reasoned that particular protein regions enriched for predicted IMMs could present a therapeutic opportunity in certain cancer types. Using HotMaps 1D, we identified clusters of residues within protein regions having statistically significant enrichment of predicted IMMs (q<0.01). These included *CIC* in low grade glioma (LGG); *NFE2L2* and *FGFR3* (Figure 7D) in bladder cancer (BLCA), *KRAS* in pancreatic adenocarcinoma (PAAD), *KIT* in testicular germ cell tumors (TGCT), *HRAS* in head and neck squamous carcinoma (HNSC); *PTEN*, *POLE* and *PPP2R1A* in uterine corpus endometrial carcinoma (UCEC) and *GNAQ* and *SF3B1* in uveal melanoma (UVM). Three genes were notable for harboring predicted immunogenic regions in more than one cancer type: *P53* in BLCA, BRCA, HNSC, LGG and UCEC, *PIK3CA* in HNSC and cervical squamous cell carcinoma (CESC) and *CTNNB1* in Liver Hepatocellular Carcinoma (LIHC) and UCEC. (Table S9d)

We explored the relationship between mutation driver status predicted by CHASMplus, and IMM status using logistic regression. The log-odds of being an IMM was significantly decreased for drivers (β=−0.66, Wald test p<2e−16), which is consistent with previous work suggesting that negative evolutionary selection eliminates MHC Class I immunogenic oncogenic mutations early in tumor development (57).

Finally, we considered whether a patient’s predicted IMM load was associated with changes in immune cell infiltrates as estimated from RNA sequencing of bulk cancer tissue. IMM load was significantly associated with increased total leukocyte fraction (β=0.75, Wald test p<2e−16) and with increased CD8+ T-cell fraction (β=5.9, Wald test p<2e−16).

These findings suggest a central role of IMMs in driving tumor immunoediting and may be informative for the interpretation of responses in the setting of immunotherapy.

## Discussion

MHCnuggets provides a flexible open-source platform for MHC-peptide binding prediction that can handle common MHC Class I and Class II alleles, as well as rare alleles of both classes. The LSTM network architecture can handle peptide sequences of arbitrary length, without shortening or splitting. The single neural network architecture requires fewer hyperparameters than more complex architectures and simplifies network training. In addition, our neural network transfer learning protocols allow for parameter sharing among allele-specific, binding affinity -and HLAp-trained networks. When trained on binding affinity data, MHCnuggets achieves comparable performance to current methods. When trained on both binding affinity and HLAp data, we demonstrate significantly improved PPV_n_ on an independent HLAp test set, with respect to other methods that use both binding affinity and HLAp data. Although PPV_n_ was lowest for the independent HLAp test set for all methods, this result is likely due to systematic differences between training HLAp data (monoallelic B-cell lines) (27) and the test data comprised of seven multi-allelic cell lines (HeLA, HTC116, JY, fibroblasts, SupB15, HCC1937, HCC1143)(24) (7), yielding a more challenging prediction problem. We attribute MHCnuggets’ improvement on the independent test set with respect to other methods to: 1) optimization of PPV_n_ in our network training protocol; and 2) our implementation of transfer learning to integrate information from binding affinity and HLAp measurements. Notably the performance of all methods is generally highest when both training and test data come from similar binding affinity experiments, but performance improvement on HLAp data is more biologically relevant (24).

We demonstrate improved scalability by comparing the runtime of MHCnuggets on 1 million peptides to comparable methods, and further by processing over 26 million expressed peptide-allele pairs across TCGA samples in under 2.3 hours. We identified 101,326 unique immunogenic missense mutations (IMMs) harbored by patients using 26 cancer types sequenced by the TCGA, based on transcriptional abundance and differential binding affinity compared to reference peptides. These results contrast with a previous report of neoantigens in TCGA patients in several respects. Rech *et al.* (45) applied a minimum expression threshold of 1 RNA sequencing read count, an IEDB-recommended combination of neoantigen predictors derived primarily from different versions of NetMHC, and IC50 threshold of 50nM to identify strong MHC binders. Their approach yielded 495,793 predicted Class I classically defined neoantigen peptides (each harboring a single immunogenic mutation) from 6,324 patients in 26 cancer types. As in our study, high variability in neoantigen burden across cancer types was observed. The striking difference between IMM and neoantigen burden in the two studies is likely due to differences in RNA expression threshold and the low false positive rate of MHCnuggets compared to IEDB-recommended tools.

Based on our conservative thresholds, IMMs were almost exclusively private to individual TCGA patients, with only 1,393 IMMs observed in more than one patient. Although more than 61% of IMMs shared by more than two patients were predicted to be driver mutations, the overall log odds of immunogenicity significantly decreased for predicted driver mutations, indicating immunogenicity might shape the driver mutation landscape. Patient IMM counts were also significantly associated with increase in total leukocyte fraction and fraction of CD8+ T-cells, suggesting that they may be relevant to immune system response to cancer.

This work has several limitations. First, our analyses are limited to missense mutations, and while these are very numerous, there is substantial evidence that somatic gene fusions, frameshift indels, splice variants etc. in tumors may also generate neoantigens. While MHCnuggets can handle peptide sequences regardless of their mutational origins, we prioritized missense mutations in this study. Next, recent work suggests that peptidal context, such as flanking sequence, its source protein and the expression level of the source protein, is informative for MHC ligand prediction (21,27). This type of information is currently only available for a limited number of HLAp data sets, which were unavailable to us for training purposes. As more well-characterized HLAp datasets become available, we will extend MHCnuggets to include these features. We did not address T-cell receptor (TCR) binding to bound peptide-MHC complexes or T-cell activation upon complex binding. While we are actively pursuing this more complex modeling problem, we believe that improved prediction of peptide binding to MHC is also therapeutically relevant (21). Finally, we are unable to directly compare performance to the MHC Class II prediction methods from the NetMHC group, except for self-reported auROC. While we are not able to do a rigorous comparison of MHCnuggets Class II prediction, our benchmark comparisons suggested that MHCnuggets was competitive with NetMHCII2.3 and that MHCnuggets Class II rare allele performance was competitive with NetMHCIIpan3.2. Generally rare allele performance estimated for each allele, regardless of MHC Class or performance metric was variable among individual alleles for both MHCnuggets and NetMHCIIpan3.2, suggesting that further work in this area is warranted.

In summary, we present MHCnuggets, an open source software package for MHC ligand prediction that improves on performance of previous methods with respect to positive predictive value by leveraging transfer learning to integrate binding affinity and HLAp data. In contrast to previous methods, it handles both MHC Class I and Class II ligand prediction and both common and rare HLA alleles, within a single framework. The utility of MHCnuggets is demonstrated with a basic pipeline for large-scale cancer patient sequencing data from TCGA, which analyzed mutation immunogenicity, shared IMMs and the relationship between mutation immunogenicity, driver potential and immune infiltrates.

## Acknowledgments

Part of this research project was conducted using computational resources at the Maryland Advanced Research Computing Center (MARCC).

