## Supplementary Information for "High-throughput prediction of MHC Class I and Class II neoantigens with MHCnuggets"

### Methods

##### Implementation

*Transformation of peptide binding affinities*. Predicted binding affinity can be transformed into a range of values well-suited for neural network learning by selecting a logarithmic base to match the weakest binding affinity of interest (1). For most benchmarks in this work, we used the standard upper limit of 50,000 nM, so that predicted binding affinity was $y = max(0,1 - log_{50k}(IC50))$. For the Bonsack et al. dataset (2), the upper limit was changed to 100,000nM because in their experiments, as described in O'Donnell et al. (3), binders were defined as peptides with IC50<100,000nM. As binding affinity was determined based on in vitro HLA binding-competition vs. a known strong binder (reported IC50 <50nM) experimental IC50 values were in µM range.

*Selection of final network weights.* To minimize overfitting, network training was stopped after 100 epochs but if the best $PPV_n$ was reached earlier, network weights from that earlier epoch were used in the final network. Notably, while we chose to optimize the networks on $PPV_n$, an alternative approach could optimize on auROC, Kendall's *tau* or Pearson's *r* correlation. For the two alleles in IEDB with the most training examples in their respective class, HLA-A*02:01 for Class I and HLA-DRB2*01 for Class II, training was stopped after 200 epochs.

*Network training.* Mean-squared error loss $L_{MSE}$ was used to train networks with continuous-valued binding affinity data and binary cross-entropy loss $L_{BCE}$ for binary HLAp data. For a dataset with *n* samples,

$$L_{MSE}(\hat{y}, y) = \frac{1}{n}\sum_{i=1}^{n}(y^{(i)} - \hat{y}^{(i)})^2$$

$$L_{BCE}(\hat{y}, y) = -\frac{1}{n}\sum_{i=1}^{n} y^{(i)}log(\hat{y}^{(i)}) + (1 - y^{(i)})log(1 - \hat{y}^{(i)})$$

All training used backpropagation with the Adam optimizer (4) and learning rate of 0.001. Regularization was performed with dropout and recurrent dropout (5) probabilities of 0.2. The number of hidden units, dropout rate, and number of training epochs was estimated by three-fold cross-validation on MHC Class I A*02:01, a common allele with a large number of experimentally characterized binding peptides.

*One-hot encoding:* Peptides were represented to the network as a series of amino acids; each amino acid was represented as a 21-dimensional smoothed, one-hot encoded vector (0.9 and 0.005 replace 1 and 0, respectively).

*Peptide padding:* MHCnuggets' architecture is capable of handling peptides of any length, but in practice a maximum length should be selected, which in this work was 15 for Class I and 30 for Class II). Peptides that are less than the maximum length are padded at the end with a character ("Z" which is not in the amino acid alphabet) until they reach the maximum length.

*Transfer Learning Protocol for binding affinity data only.* We used transfer learning to improve network learning for MHC alleles with limited characterized peptides available for training. We first trained base allele-specific networks for Class I and Class II, using alleles with the most training examples in IEDB (HLA-A*02:01 for Class I and HLA-DRB2*01 for Class II). For all other alleles, the final weights of the base network for its respective class were used to initialize network training, and then an allele-specific network was trained for each allele. Next, we assessed prediction performance of each allele-specific network on the training examples for each of the alleles. For each allele, if the network that performed best was not the HLA-A*02:01 network (for Class I alleles) or HLA-DRB1*01:01 network (for Class II alleles), we did a second round of training, with the best performing network's weights used in the initialization step.

*Transfer Learning Protocol for binding affinity and HLAp data.* To integrate HLAp data into the Class I networks we initially trained each network with binding affinity data as described above, transferred the final weights to a new network, and then continued training with the HLAp data as positive examples augmented with random peptide decoys as negative examples.

**Dataset collection and curation**

Table S1. Data sources used in training and benchmarking. BA=binding affinity. HLAp=peptide elution/mass spectrometry. LOMO=leave one molecule out cross-validation. Only alleles with >30 characterized peptides were included. IEDB=curated version 2018. Common allele=>30 peptides with characterized binding information, rare allele= <30 characterized peptides, mono-allelic = engineered cells that express a single MHC allele, multi-allelic = cells that express multiple MHC alleles. Abelin et al. (6), Bonsack et al. (2), Kim et al. (7) Bassani-Sternberg et al. (8) Trolle et al. (9), Jensen et al. (10), IEDB (11).

| MHC Class | Common or rare alleles | Training Sets | | Benchmarks | |
|---|---|---|---|---|---|
| | | **Datasets** | **Description** | **Datasets** | **Description** |
| Class I | Common | *IEDB* | BA, 241,553 peptides for 217 alleles | *Bonsack et al* | BA, Tested on 475 peptides for 7 alleles |
| | | *Abelin et al* | HLAp, 23,651 peptides for 16 alleles, mono-allelic | *Kim et al* | BA, Trained on 53 alleles (BD2009), tested on 53 alleles (BLIND) |
| | | | | *BST* | HLAp, 23,971 hits for 6 alleles .from Bassani-Sternberg 2017 and Trolle et al. plus random peptide decoys. |
| | | | | Bassani-Sternberg et al 2017 | HLAp, 22,598 hits for 26 alleles, multi-allelic Included in BST. |
| | | | | Trolle et al | HLAp, 15,524 hits for 5 alleles, mono-allelic Included in BST. |
| | | | | Random peptide decoys from human proteome | 23,947,029 random decoy peptides generated from the human proteome (Included in BST). |
| Class II | Common | *IEDB* | BA, 96,211 peptides for 135 alleles | *Jensen et al* | BA, Five-fold cross-validation on 27 alleles. |
| Class I | Rare | NA | NA | *IEDB Pseudo-Rare Alleles* | BA, LOMO on 20 alleles, alleles with training samples between 30 and 100 |
| Class II | Rare | NA | NA | *Jensen et al* | BA, LOMO on 27 alleles |

Data sources for network training and testing, TCGA somatic mutations, TCGA tumor gene expression and haplotype calling are shown in Table S1 and Table S2. A curated version of the IEDB database 2018 (11) and the sixteen Class I mono-allelic B-cell line immunopeptidomes (6) was provided by Tim O'Donnell (https://data.mendeley.com/datasets/8pz43nvvxh/2), binding affinity assays of HPV-derived peptides were provided by Maria Bonsack and Angelika Riemer (2), BST = immunopeptidomes from six cell lines with multi-allelic MHCs (26 MHC Class I alleles) (8) and from soluble HLA(sHLA)-transfected HeLa cells separated by allele (4 MHC Class I alleles) (9). Decoy random peptides were sampled from the human proteome.

**Training Data Sets**

The networks were trained with data from a curated version of the Immune Epitope database (IEDB 2018) (11), containing chemical binding affinity measurements for 241,553 peptide-allele pairs covering 217 Class I alleles and 96,211 peptide-allele pairs covering 135 Class II alleles. Additional training data consisted of 16 Class I mono-allelic B-cell line immunopeptidomes (6). The immunopeptidome data is limited to HLAp binders and was supplemented by decoy random peptides sampled from the human proteome (https://data.mendeley.com/datasets/8pz43nvvxh/2).

**Benchmark Data Sets**

*Kim et al:* This benchmark contained 53 MHC Class I alleles and 137,654 IC50 measurements published prior to 2009 (training set) and 53 unique MHC Class I alleles with 26,888 IC50 measurements, published from 2009-2013 (test set). Three alleles (HLA-B*27:03, HLA-B*38:01, HLA-B*08:03) did not contain sufficient training data, and two alleles (HLA-A*46:01, HLA-B*27:03) did not contain any peptides defined as binders in this work (IC50<500nM). Therefore, a total of four alleles (HLA-A*46:01,HLA-B*27:03, HLA-B*38:01, and HLA-B*08:03) were dropped from the analysis. All peptides in this benchmark set consisted of 8-11 amino acid residues.

*Bonsack et al.* This dataset contains 475 synthetic peptides derived from model protein sequences HPV16 E6 and E7 tested for binding to 7 alleles (HLA-A*01:01, HLA-A*02:01, HLA-A*03:01, HLA-A*11:01, HLA-A*24:02, HLA-B*07:02 and HLA-B*15:01). Each peptide was tested in competition-based cellular binding assays with a known high-affinity fluorescein-labeled reference peptide. EBV-transformed B-lymphoblastic cells were stripped of their naturally-bound peptides and mixed with serially diluted test peptides and 150 nM of reference peptide. Each synthetic peptide was tested at 8 different concentrations ranging from 780 nM to 100,000 nM. Mixture fluorescence at each synthetic peptide concentration was measured with flow cytometry, and a non-linear regression analysis was used to find the test peptide concentration that inhibited 50% of the reference peptide binding (IC50). Peptides were classified as binders (IC50 <= 100,000 nM) or nonbinders (IC50 > 100,000 nM). Peptides in this independent benchmark set do not have IEDB entries.

*Bassani-Sternberg et al. 2017* (8) This dataset contains 22,598 unique peptides eluted from 6 cell lines with multi-allelic MHCs. Out of the total 6 cell lines, a total of 26 alleles were reported. For each multi-allelic cell line, peptide/MHC pairs were found through deconvolution, following the protocol described by (6), with the difference that we used MHCnuggets rather than NetMHCpan2.8 (12) to predict IC50 values for each peptide-MHC pair. For each cell line, each peptide was initially assigned as a binder to all expressed alleles. Then, for each allele, we filtered out any peptide predicted to bind with IC50>1000nM to that allele, and with IC50<150nM to any other allele. Peptides found for 6 alleles (HLA-A*01:01, HLA-A*02:01, HLA-A*03:01, HLA-A*24:02, HLA-A*31:01, HLA-B*51:01), were selected for allele-specific prediction testing. Trained networks were available for these alleles from all the methods that we compared.

*Trolle et al.* This dataset contains 15,524 unique peptides eluted from soluble HLA(sHLA) transfected HeLa cells, a process that allowed for separating binding peptides to a single MHC allele. This dataset reports peptides for 5 MHC alleles. Peptides found for 4 alleles (HLA-A*01:01, HLA-A*02:01, HLA-A24*02, HLA-B*51:01) were selected for testing. Peptide lengths in this dataset range from 8-15 amino acid residues.

*BST.* This benchmark consists of 23,971 HLAp hits for 6 alleles, from Bassani-Sternberg et al. 2017 and Trolle et al. plus 23,947,029 random decoy peptides sampled from the human proteome. Any peptides found to overlap with the training HLAp data (Abelin et al.) were removed.

*Jensen et al.* This benchmark was designed to assess both allele-specific and rare MHC Class II binding affinity predictors. Allele-specific prediction was tested with a five-fold cross validation experiment on peptides found in IEDB in 2016 but not 2013. Rare allele predictions were tested with the LOMO protocol.

*IEDB Class I rare alleles.* This dataset was designed to apply the LOMO protocol to Class I alleles. It included 20 "pseudo-rare" alleles with 30-100 binding affinity peptide measurements in IEDB.

**<u>Performance metrics</u>**

We calculated positive predictive value with respect to the top-ranked *n* peptides, where *n* is the number of true binders in the ranked list, denoted as $PPV_n$.
PPV = NTP / (NTP+NFP), where NTP=number of true positives and NFP=number of false positives. We calculated PPV with respect to the top-ranked n peptides, where n is the number

of true binders in the ranked list, denoted as $PPV_n$. For the BST benchmark, we also calculated PPV over the top 50 and 500 ranked peptides.

**Runtime analysis**

To assess the speed and scalability of the tested methods, we selected one million peptides sampled from the Abelin et al. dataset (6) for Class I alleles, and one million peptides sampled from the IEDB (curated dataset 2018 (13)) for Class II alleles. Sampling was done with replacement. For each method listed in Figure 1A, networks for three Class I MHC alleles (HLA-A*02:01, HLA-A*02:07, HLA-A*01:01) and three Class II MHC alleles (HLA-DRB1*01:01, HLA-DRB1*11:01, HLA-DRB1*04:01) were used to predict binding over a range of input sample sizes ($10^2$, $10^3$, $10^4$, $10^5$, $10^6$ ). All methods were run on a single GPU compute node (one NVIDIA TESLA K80 GPU plus six 2.50GHz Intel Xeon E5-2680v3 CPUs, 20GB memory).

**TCGA analysis pipeline**

*MC3 mutation filtering:* MC3 TCGA somatic mutation calls were filtered for missense mutations.

Cancer types in the TCGA are abbreviated as follows:

| | |
|---|---|
| ACC | Adrenocortical carcinoma |
| BLCA | Bladder Urothelial Carcinoma |
| BRCA | Breast invasive carcinoma |
| CESC | Cervical squamous cell carcinoma and endocervical adenocarcinoma |
| CHOL | Cholangiocarcinoma |
| COAD | Colon adenocarcinoma |
| GBM | Glioblastoma multiforme |
| HNSC | Head and Neck squamous cell carcinoma |
| KICH | Kidney Chromophobe |
| KIRC | Kidney renal clear cell carcinoma |
| KIRP | Kidney renal papillary cell carcinoma |
| LGG | Brain Lower Grade Glioma |
| LIHC | Liver hepatocellular carcinoma |
| LUAD | Lung adenocarcinoma |
| LUSC | Lung squamous cell carcinoma |
| PAAD | Pancreatic adenocarcinoma |
| PCPG | Pheochromocytoma and Paraganglioma |
| PRAD | Prostate adenocarcinoma |
| READ | Rectum adenocarcinoma |
| SARC | Sarcoma |

| | |
|---|---|
| TGCT | Testicular Germ Cell Tumors |
| THCA | Thyroid carcinoma |
| THYM | Thymoma |
| UCEC | Uterine Corpus Endometrial Carcinoma |
| UCS | Uterine Carcinosarcoma |
| UVM | Uveal Melanoma |

*Regression models:* We applied two univariate Poisson regression models. In the first model, each patient's predicted immunogenic missense mutation load was the response variable and the independent variable X was the total leukocyte fraction. The fitted coefficient $\beta = 0.75$ (p<2e-16, Wald test) indicated that increased predicted IMM load was significantly associated with increased leukocyte fraction in a patient's cancer. In a second model, X was the proportion of CD8+ T cells inferred by CIBERSORT (14). The fitted coefficient $\beta = 5.9$ (p<2e-16, Wald test) indicated that increased predicted IMM load was associated with increased tumor-infiltrating CD8+ T cells. Total lymphocyte and (Aggregate3) CD8+ T cell fractions were estimated in Thorsson *et al.* (15).

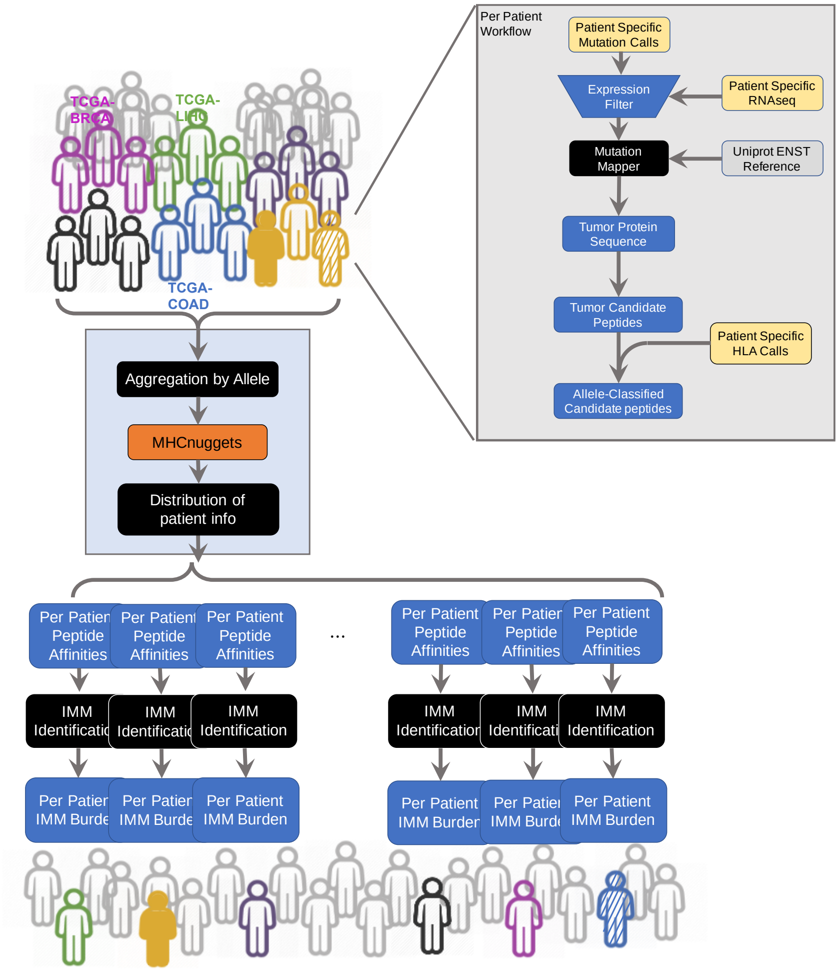


**Figure S1. Flow chart describing the neoantigen prediction pipeline applied to TCGA.** TCGA mutation calls were filtered by transcript expression for each patient. Mutations were mapped to reference transcripts and protein sequences. Peptides of length 8-11 were extracted from patients' mutated protein sequences. Candidate peptides for each mutation were selected by differential binding affinity to up to six possible Class I alleles from each patient (Methods). TCGA samples were processed with an "hourglass" design. Peptides were aggregated by allele, and MHCnuggets predicted binding affinities were calculated across all patients for each allele. Peptides that passed all filters were considered candidate neoantigens and were re-assigned to the originating patient. Predicted IMMs are defined to be the somatic missense mutations that generate candidate neoantigen peptides. Predicted IMM load was computed for each patient and in aggregate for each cancer type. IMM=immunogenic missense mutation.
